# langevitour: smooth interactive touring of high dimensions, demonstrated with scRNA-Seq data

**DOI:** 10.1101/2022.08.24.505207

**Authors:** Paul Harrison

**Affiliations:** Monash Bioinformatics Platform, Monash University, 15 Innovation Walk, Monash University, Clayton Campus, Clayton VIC, Australia, 3800

## Abstract

langevitour displays interactive animated 2D projections of high-dimensional datasets. Langevin Dynamics is used to produce a smooth path of projections. Projections are initially explored at random. A “guide” can be activated to look for an informative projection, or variables can be manually positioned. After a projection of particular interest has been found, continuing small motions provide a channel of visual information not present in a static scatter-plot. langevitour is implemented in Javascript, allowing for a high frame rate and responsive interaction, and can be used directly from the R environment or embedded in HTML documents produced using R. The widget is demonstrated using single-cell RNA sequencing (scRNA-Seq) data. langevitour’s linear projections provide a less distorted view of this data than commonly used non-linear dimensionality reductions such as UMAP.

## Introduction

Understanding high-dimensional data is difficult. As a motivating example, single cell RNA-sequencing (scRNA-Seq) typically measures the expression levels of thousands of genes in tens of thousands of cells. Thinking of cells as points in a space where the dimensions are expression levels of each gene, there is a complex high-dimensional geometry due to differences between cell types, variation in expression within cell types, cell developmental trajectories, and responses to treatments. Most of this variation can be summarized down to tens of dimensions using techniques such as Principal Components Analysis (PCA), but even a ten-dimensional space is difficult to comprehend.

One way to explore high-dimensional data is using a “tour.” A tour is a sequence of projections of the dataset, most commonly into two dimensions. A Grand Tour (Asimov, 1985) is a tour that will eventually visit as close as we like to every possible projection of the data, typically using a sequence of random projections. A Guided Tour on the other hand seeks an “interesting” projection by moving toward the maximum of some index function (Cook et al., 1995). The sequence of projections is animated, with smooth interpolation between each successive pair of projections. The software XGobi and GGobi (Swayne et al., 1998) provide an interactive graphical application incorporating tours for exploring high-dimensional data. The more recent R package **tourr** (Wickham et al., 2011) provides a framework for creating and displaying tours in the R language. Displaying animations directly in R usually does not achieve a high frame-rate. It is also not possible to interact with the display as was possible with GGobi. To get around these problem, a recent R package called **detourr** (Hart and Wang, 2022) computes a tour path in R using **tourr** and then displays it using a Javascript widget (using **htmlwidgets**) (Vaidyanathan et al., 2021). The widget then provides a high frame-rate display and interactive features. However the projection path itself can not be modified interactively.

This article introduces a new R package, **langevitour. langevitour** differs from previous tour software by using Langevin Dynamics, a method from physics, to produce a continuous path of projections. This path can be directly used for animation, doing away with the need to interpolate between distinct projections to animate the tour. The package is **htmlwidgets**-based, with interaction, calculations, and animation performed in Javascript. This allows the projection to be controlled interactively so that the user may switch between Grand and Guided Tours, while also interactively focussing in on particular dimensions of interest.

Langevin Dynamics is described mathematically in the methods section, but I will outline its important features here using two physical examples. First, consider modelling the position and velocity of a large particle over time. The large particle is continuously jostled by many small particles. This is the original Brownian motion scenario studied by Langevin in 1908 (see translation by Lemons and Gythiel, 1997). Rather than modelling every particle, Langevin Dynamics simulates the jostling as small random forces. Langevin’s model includes these random forces and damping of momentum, and we can also add a force field acting on the particle. The particle explores the space it is in, and the force field may cause the particle to spend more time near certain locations.

**langevitour** applies Langevin Dynamics not to a particle’s position but rather to an orthonormal projection matrix. As a second physical example, imagine a two-dimensional disk in a high-dimensional space. The disk represents a projection plane for a high-dimensional dataset. It has a fixed center but is able to rotate freely. Tiny unseen particles continuously jostle the disk, causing it to spin first one way and then another. The motion of the disk provides the path for a Grand Tour of the dataset. A force field may also draw it toward particular orientations. The force field is specified using a potential energy function and is used to seek interesting projections of the data, similar to the index functions used in previous tour software, providing a Guided Tour.

## Results

### Palmer Penguins example

The R data package **palmerpenguins** (Horst et al., 2020) provides body measurements of penguins of three different species from the Palmer Archipelago, Antarctica. The **langevitour** visualization is shown in Figure 2. R code to produce this figure is:

**Figure 1:**
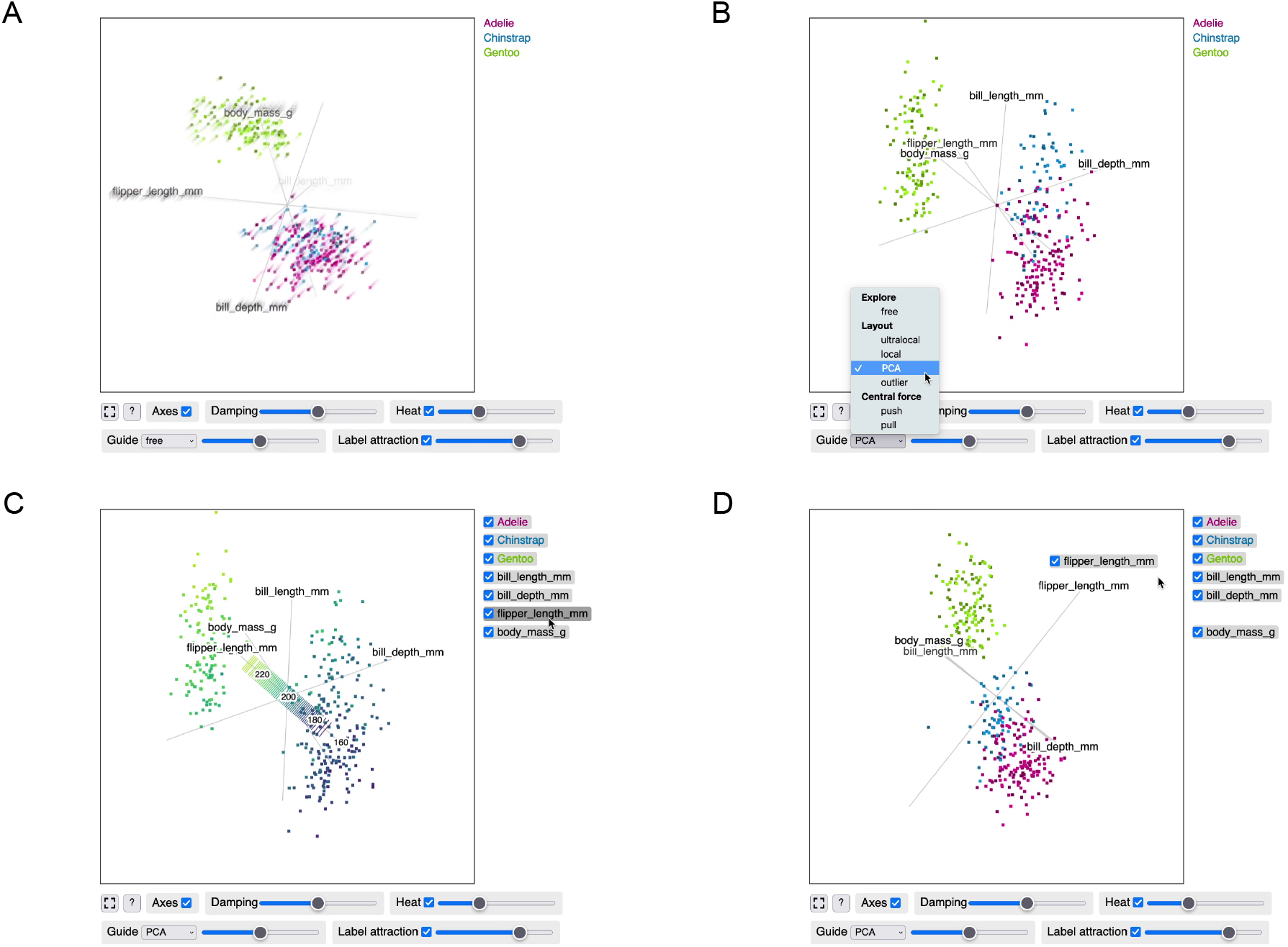
**palmerpenguins** data visualized using **langevitour**. An interactive version of this figure is available in the supplemental file figures-page.html. (A) Spinning at random. (B) Activating a guide. (C) Mousing over a label to color by a variable. (D) Dragging a label onto the plot to concentrate on a particular variable.

**Figure 2:**
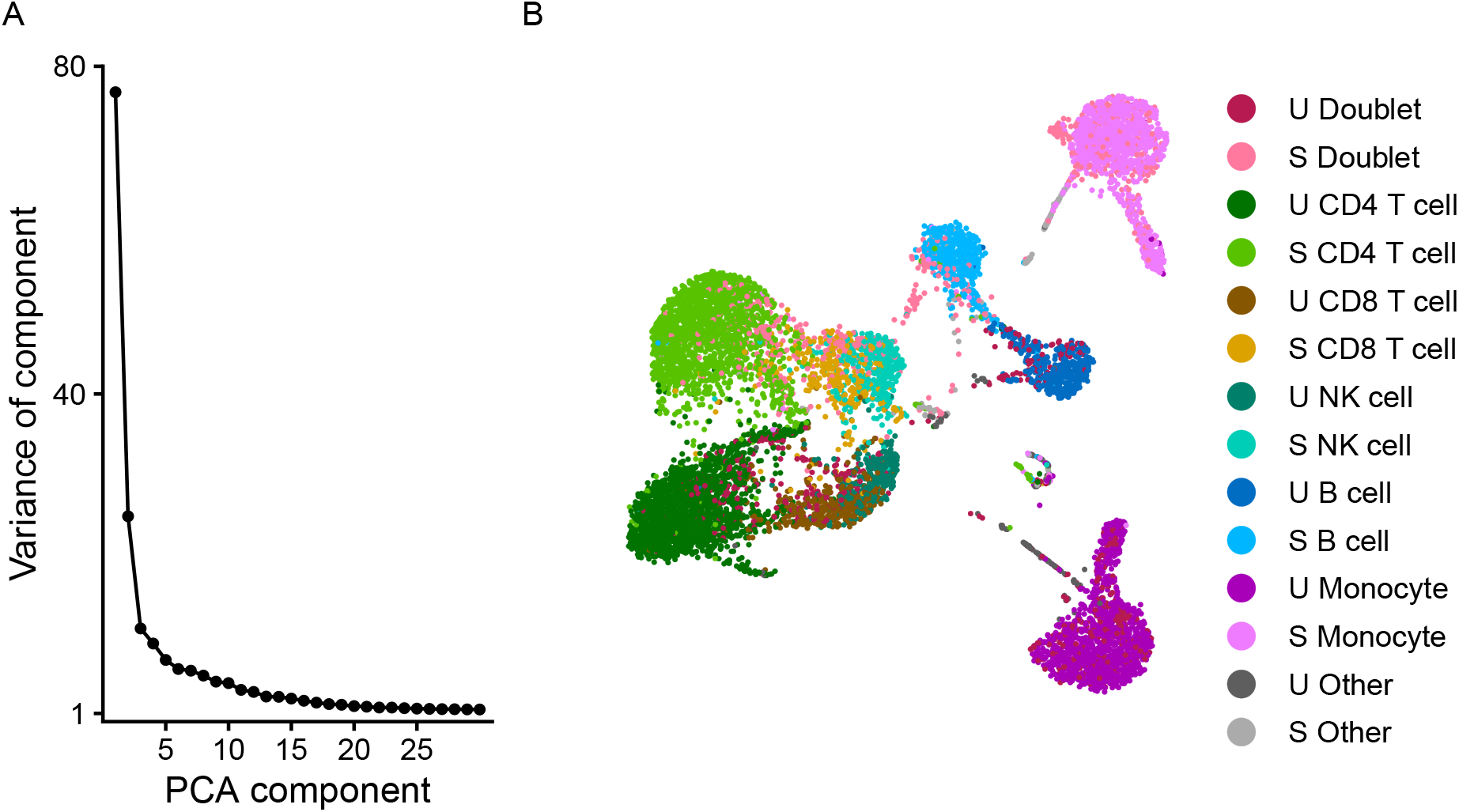
scRNA-Seq analysis using Seurat. (A) Scree plot. (B) UMAP layout based on the cell scores of the first 10 PCs. U are unstimulated and S are stimulated cells.

~~~
library(langevitour)
library(palmerpenguins)
completePenguins <-na.omit(penguins[,c(1,3,4,5,6)
scale <-apply(completePenguins[,-1], 2, sd)*4
langevitour(completePenguins[,-1], completePenguins$species,
      scale=scale, pointSize=2)
~~~

The **langevitour** interface allows:

- Setting a “guide.” This causes **langevitour** to pursue projections in the vicinity of the minimum of an energy function. For example the PCA guide seeks projections with large variance in both the x and y directions.
- Hiding particular groups by unchecking their checkbox, in order to focus on other groups. For example by hiding Gentoo penguins we can focus on the difference between Adelie and Chinstrap penguins. The guide is also only applied to the visible groups.
- Hiding particular axes by unchecking their checkbox. For example without bill length Adelie and Chinstrap penguins can no longer be distinguished.
- Dragging labels on to the plot to concentrate on particular axes or try to separate out a particular group. The projection may not exactly match the label positions, since it must still be orthonormal.
- Adjusting damping. High damping produces jerky Brownian motion. Less damping produces smoother less random motion. The fastest way to thoroughly explore the space of projections is an intermediate damping level.
- Adjusting heat. More heat makes the projection move faster, and also strays further from the optimum projection defined by a guide or any labels that have been dragged on to the plot.
- Mousing over a group label. The group is highlighted.
- Mousing over an axis label. A scale and rug are displayed, and points are colored according to their position on that axis.

A systematic way to examine a dataset is to uncheck all but two or three axes at a time. With three axes checked, the eye interprets the display as three-dimensional.

If a particular interesting projection is found using **langevitour** it can be brought back in to R by pressing the “?” button and copying and pasting R code that is shown. The “?” button also shows a JSON record of the current settings of the widget, including form inputs and label positions. All or some of these settings can be specified to a future call to langevitour(), or applied to a running widget using Javascript code, for example when a button is pressed in an HTML document.

### scRNA-Seq example

A dataset by Kang et al. (2018) demonstrates many of the complex high-dimensional features found in scRNA-Seq data. In this dataset, peripheral blood mononuclear cells (PBMCs) from eight patients with lupus were pooled. PBMCs are cells from the immune system that circulate in blood, including monocytes, B cells, T cells, and natural killer (NK) cells. These cells were then stimulated with a cytokine, recombinant interferon beta, causing a change in the gene expression of the cells. The dataset contains a sample of unstimulated cells (U), and a separate sample of stimulated cells (S).

Kang et al. (2018) have made available count data from this experiment, as well as their annotation of each cell type. Slightly simplified annotations are shown in this article. For each of the two samples, there is a matrix giving the number of molecules of RNA associated with each gene within each cell. This count data was processed using **Seurat** (Hoffman, 2022) to obtain Principal Components (PCs) of the normalized and log transformed counts (details in the methods section). For each PC, there is a score for each cell and a loading for each gene.

### Cell scores

Results from analysis with **Seurat** are shown in Figure 2. There is no clear elbow in the scree plot. The top 10 PCs will be used simply as a manageable number to interact with. Common practice is to visualize cells using a 2D UMAP layout computed from the PCs, as shown in Figure 2B, to see what clusters exist in the data and try to understand the relationships between them.

UMAP (McInnes et al., 2018) is a non-linear dimensionality reduction technique. Ironically, UMAP may give a curvy biological appearance to structures that are linear in the original data! Problems with UMAP are discussed by Coenen and Pearce (2019), and are similar to the problems with t-SNE (Wattenberg et al., 2016), an earlier method that UMAP has largely supplanted for scRNA-Seq analysis. Problems include that UMAP may arbitrarily change the distances between clusters, and that by design UMAP will hide whether clusters are more or less spread-out.

Figure 3 shows the **langevitour** visualization of the cell scores. Components have been varimax rotated to improve interpretability (see methods section). To allow a smooth frame-rate, 10,000 cells were sampled out of 29,065 total cells available. With **langevitour**, we only ever see linear projections of the data. Straight lines remain straight and parallelograms remain parallelograms. Distances may be decreased but will not be increased by the projection.

**Figure 3:**
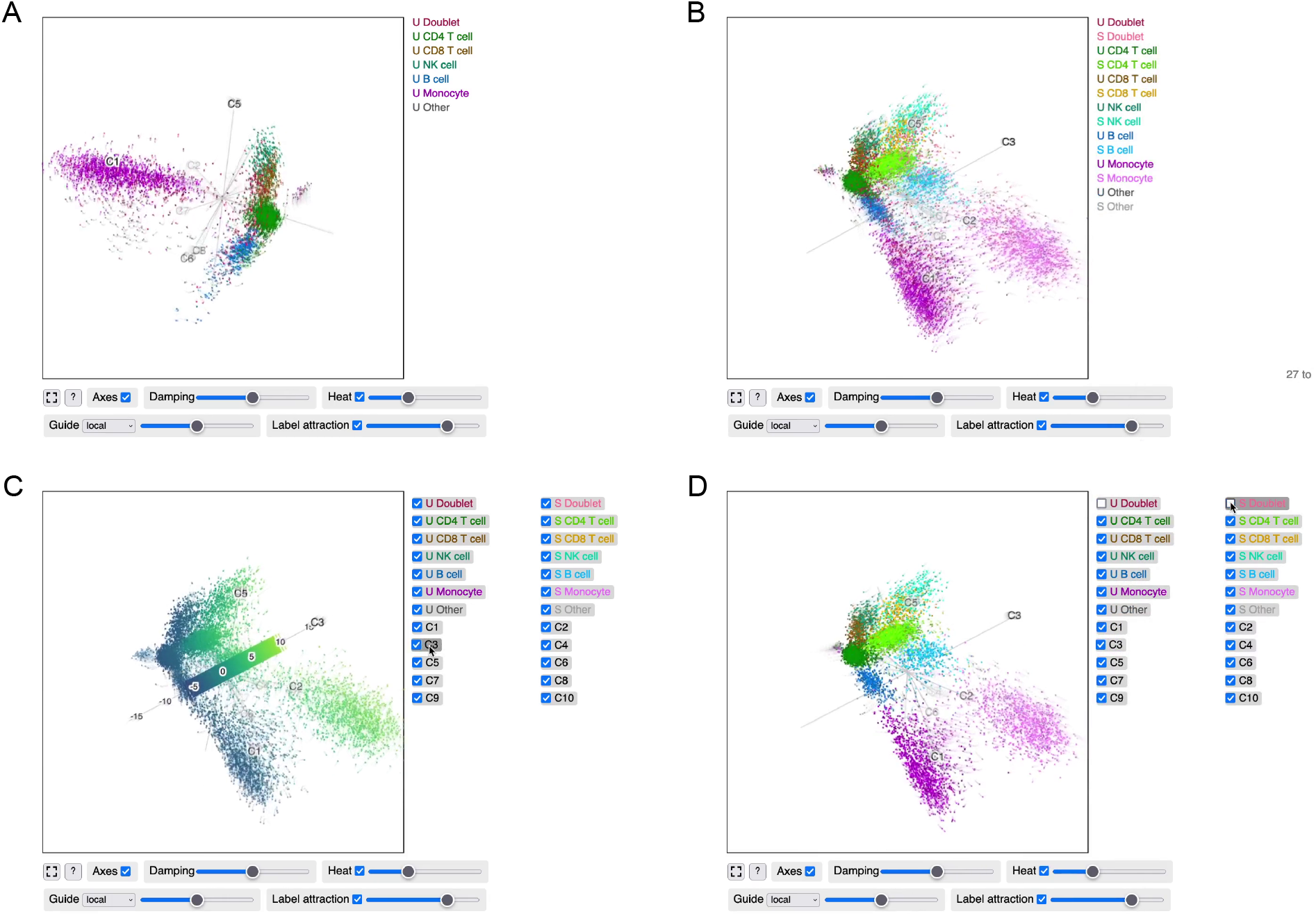
scRNA-Seq cell scores visualized using langevitour. An interactive version of this figure is available in the supplemental file figures-page.html. The “local” guide is active. (A) Unstimulated cells. (B) All cells. (C) Colored by component 3. (D) Doublets hidden.

This is a moderately complex dataset. It may be helpful to initially hide the stimulated cells and activate the “local” guide so as to see the relationships between unstimulated PBMC cells. The guided layout may not be able to perfectly separate all the clusters as UMAP can, but small relative motions make clear where clusters are overlapping by accident rather than real proximity. For example B cells and T cells are distinct.

Again, scales can be seen by mousing over axes. The scale for each component is meaningful, representing distance along a certain direction in scaled gene expression space. The direction is specified by the gene loadings, which are examined shortly.

Single cell sequencing produces a small proportion of doublets, where two cells end up in a single micro-droplet and appear in the final data as a single cell. A nice feature of this dataset is that doublets containing cells from two different individuals can be identified with certainty due to genetic differences between the individuals. Further doublets from the same individual have been imputed based on similarity to doublets between individuals. Mouse over the “U Doublet” and “S Doublet” labels to highlight doublets. Doublets located between two clusters may be interpreted as a pair of cells with different cell types. Hiding the doublets makes the clusters more distinct.

Comparing now to the UMAP layout (Figure 2B). In the linear view provided by langevitour, the monocytes can be seen to be more spread out than other cell clusters. This isn’t visible in UMAP, which as a deliberate feature erases differences in scale. The whiskers extending from various clusters in the UMAP correspond to components at right angles to other components in the data, i.e. a subset of cells in which certain genes are active. For example the thin whiskers extending from the unstimulated and stimulated monocytes actually extend in the same direction, along C7, but in the UMAP they extend in different directions. Doublets in the UMAP tend to form clumps near the edges of clusters or between clusters. In the linear view of the data they are spread out between clusters.

### Denoising

The fuzziness of the clusters in Figure 3 is an honest depiction of the data. However to interpret the geometry of the data, it may be helpful to reduce the amount of noise. We would like to do so with minimal distortion. My suggested method, implemented in the function knnDenoise(), is to average each cell with its 30 nearest neighbors and their 30 nearest neighbors. A comparison of the original and denoised positions is shown in Figure 4 and the full result is shown in Figure 5.

**Figure 4:**
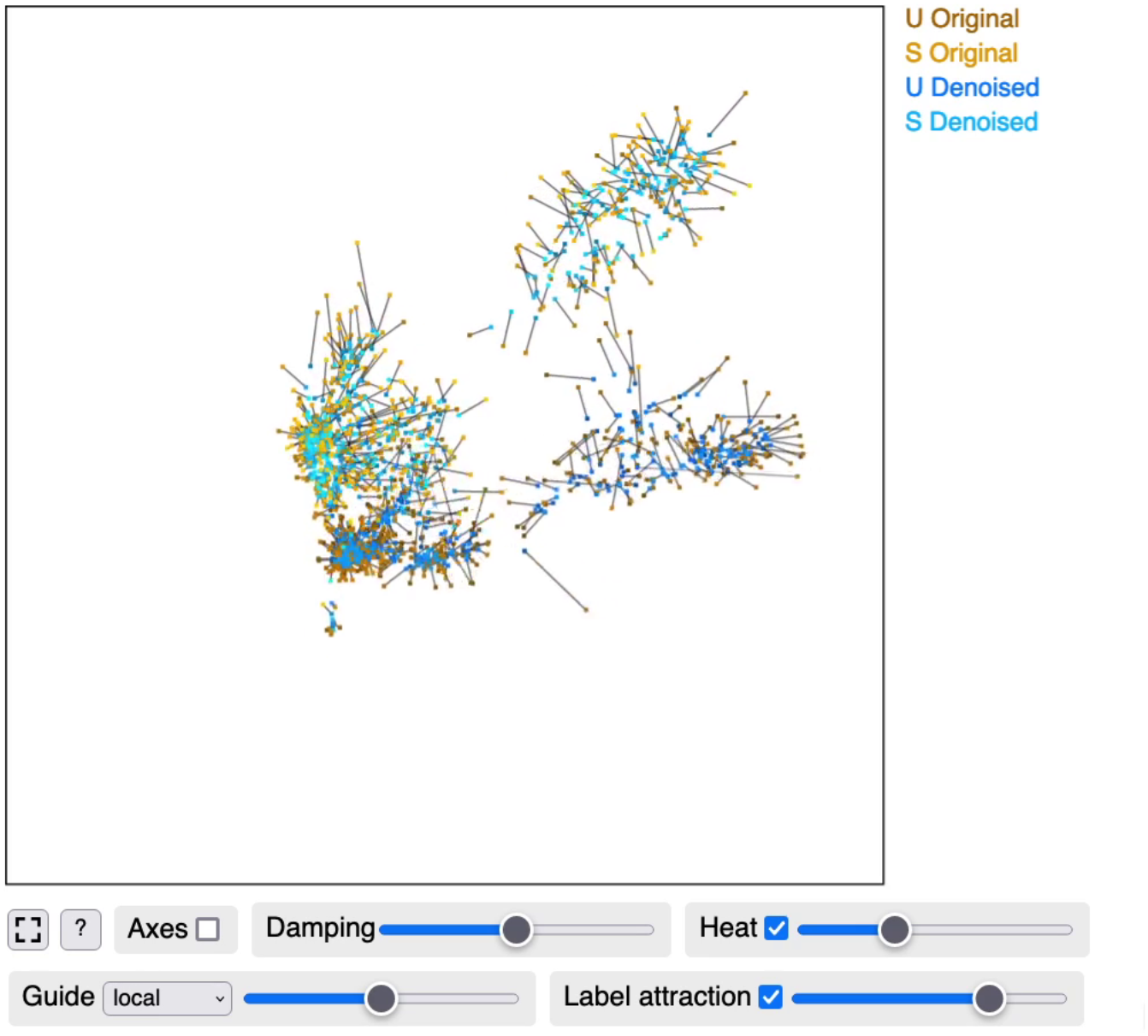
Noise reduction of scRNA-Seq cell scores. Original and denoised positions of 1,000 cells are shown, joined by lines. An interactive version of this figure is available in the supplemental file figures-page.html.

**Figure 5:**
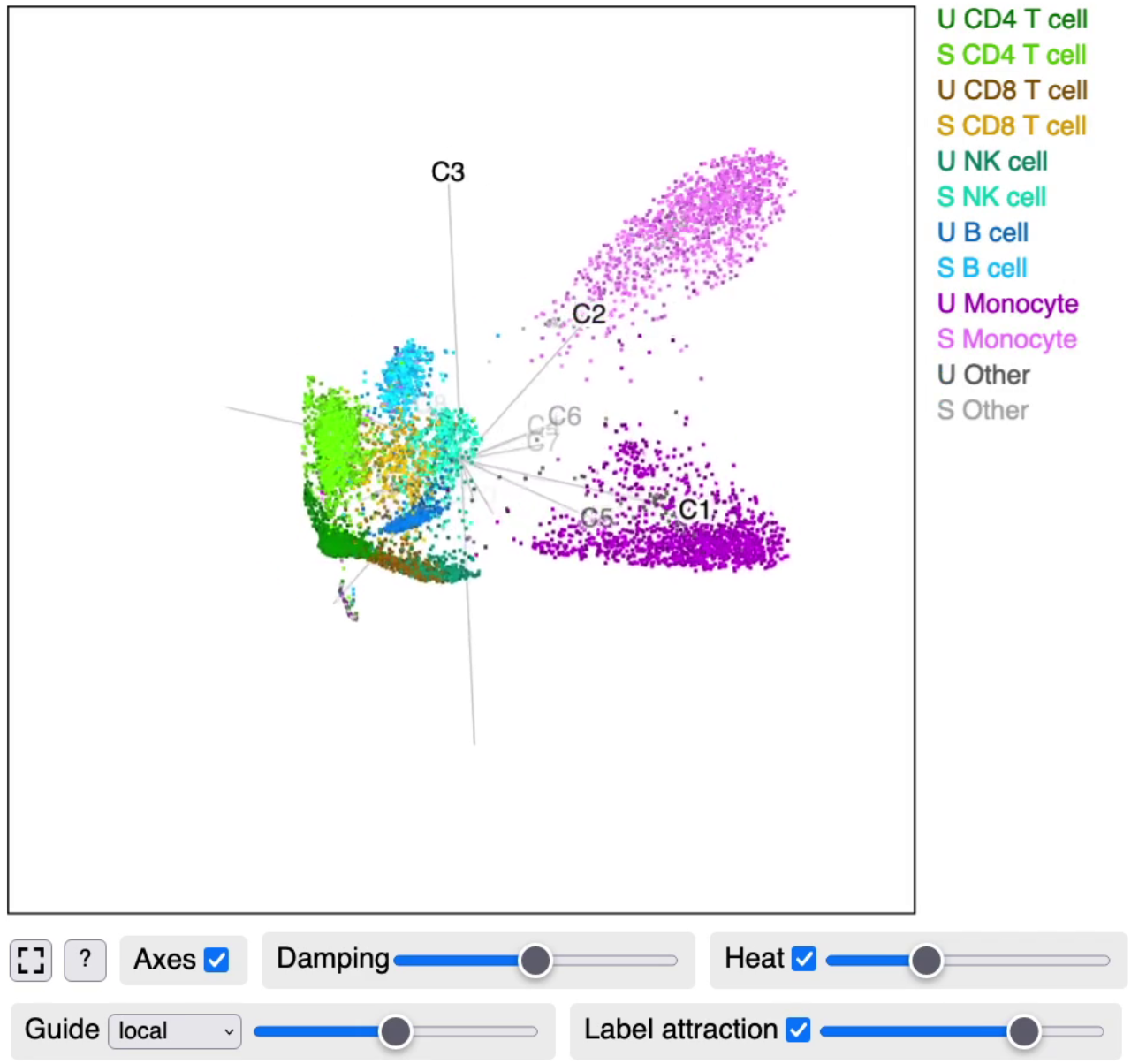
Noise reduced scRNA-Seq cell scores.

Doublets still lie between clusters in this denoised version. Compare this to UMAP layout, where the doublets tend to be attached to one or other cluster. Hiding the doublets again makes clusters cleaner.

Components associated with particular cell types can be found. C1 for monocytes, C5 for CD8 T cells and NK cells, and C6 for B cells. The response to the cytokine is mostly contained in C3 with a further monocyte specific response in C2.

### Gene loadings

The gene loadings may also be examined, as shown in Figure 6. Each component of the loadings is a unit vector.

**Figure 6:**
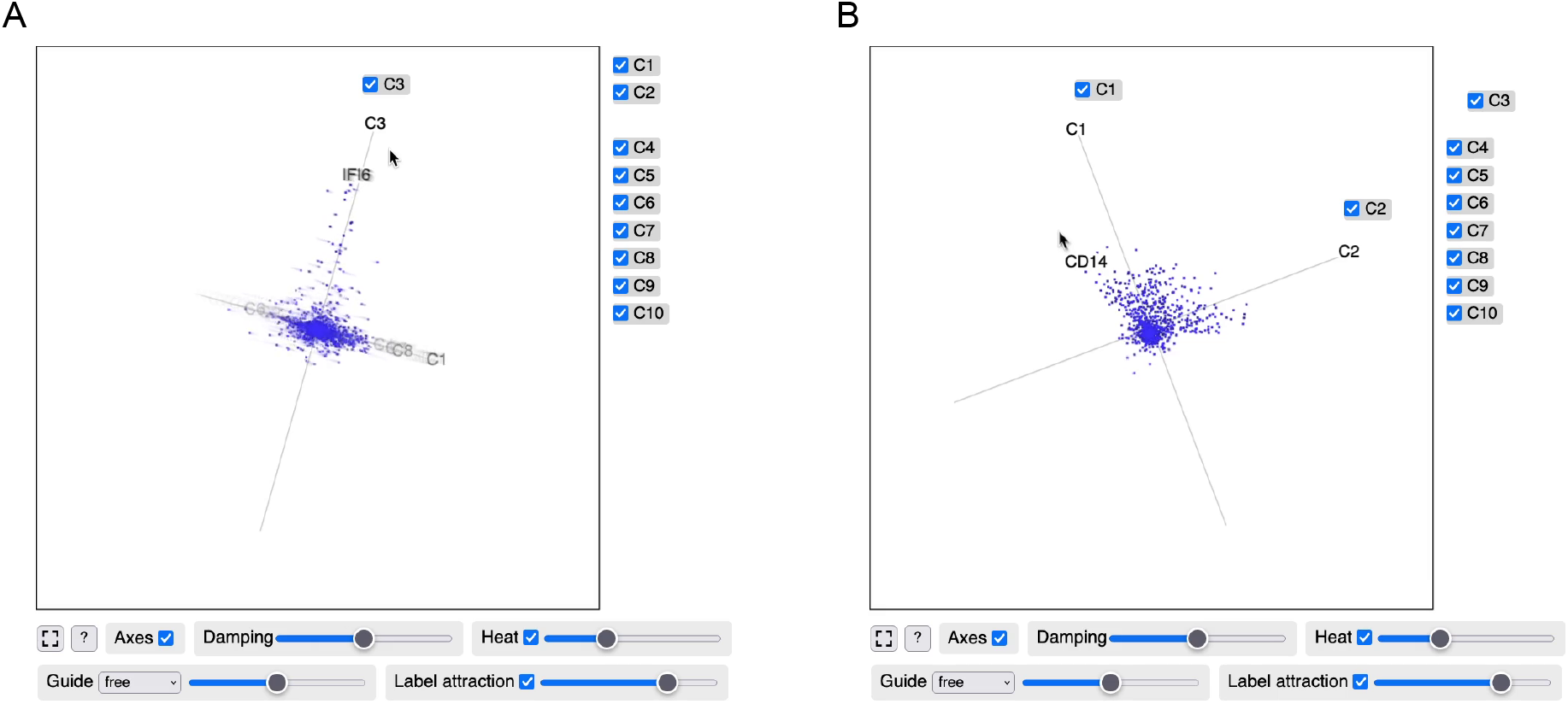
scRNA-Seq gene loadings visualized using langevitour. An interactive version of this figure is available in the supplemental file figures-page.html. (A) Genes involved in component 3. (B) Genes involved in components 1 and 2.

It can be seen that biological phenomena such as different cell types (C5, C6) or response to the cytokine (C3) often involve distinct sets of genes. Furthermore, cell types are defined more by the activation of genes than the deactivation of genes. Varimax rotation was able to align these distinct sets of genes into specific components. An exception is genes used by monocytes and the monocyte response to stimulation (C1 and C2). These two components show a broad fan of genes, which can be interpreted as the genes involved in being a monocyte also being involved to varying degrees in the monocyte response to stimulation.

Mouse near points to see the specific genes they represent.

## Methods

Say we have *n* × *p*-dimensional data points **y**_*i*_. A × 2 *p* projection matrix from *p* to 2 dimensions will be denoted **X**. A projected point is then **Xy**_*i*_.

The two rows of **X** are called a 2-frame. These rows must be be unit vectors and orthogonal to each other. The set of all 2-frames in *p* dimensions is a Stiefel manifold.

It will often be necessary to consider all of the elements of the projection matrix concatenated together into a single vector (“melted”), which will be denoted **x**.

### Langevin Dynamics overview

Projections are generated using a numerical simulation of Langevin Dynamics, but with projections constrained to lie on the Stiefel manifold. This section *briefly* summarizes Langevin Dynamics. The next section will describe the numerical simulation method and how the constraint is applied.

The description of Langevin Dynamics given here follows Leimkuhler and Matthews (2015), but for simplicity I set the Boltzmann constant to 1 and all masses to 1. We define a system having a position vector **x** and a velocity vector **v**. We will need to specify a temperature *T*, a damping rate *γ*, and a potential energy function *U*(**x**). The behavior of the system is then defined by a pair of Stochastic Differential Equations (SDEs):

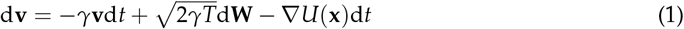

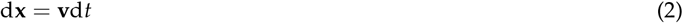

Here **W** is a vector of Wiener processes. For any positive time-step Δ*t*:

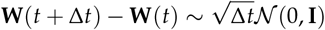

The total energy of the system, kinetic energy plus potential energy, is called the Hamiltonian:

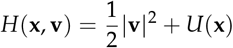

In Equation (1) in a physical system, the first two terms would describe exchange of kinetic energy with the surrounding environment. In the first term kinetic energy is lost (damping), while the second term adds randomness to the velocity, tending to increase the kinetic energy. The third term applies acceleration according to the gradient of the potential energy function.

If we were to set *γ* = 0, we would be doing Hamiltonian Dynamics and the total energy of the system would remain constant. With *γ* > 0 the total energy can fluctuate, and in the long run the process is ergodic (Leimkuhler and Matthews, 2015, in section 6.4.4), producing samples with the Gibbs-Boltzmann probability density:

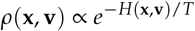

From this density it can be seen that each component of the velocity is normally distributed with variance *T* and that the position has probability density

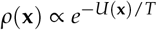

The potential energy function completely controls the distribution of positions being produced, providing a great deal of freedom. Here, we will use this to craft suitable potential energy functions to allow the user to control the projections being explored.

### Langevin Tour numerical simulation

We are to generate a sequence of animation frames *i* = 1, 2, …, each with a projection matrix **X**_*i*_, also called **x**_*i*_ when viewed as a vector. Each frame will also have a velocity vector **v**_*i*_. Call the time-step from frame *i* − 1 to frame *i* Δ*t*_*i*_.

The Position Based Dynamics method (PBD, Müller et al., 2007) is used to perform the numerical simulation while constraining the system to produce orthonormal projection matrices. PBD is simple to implement and emphasizes stability over accuracy when enforcing constraints, which is appropriate and adequate for this application. Using PBD, in each iteration we will:

1. Update the velocity.
2. Update the position based on the velocity.
3. Fix the updated position to be an orthonormal projection matrix.
4. Fix the velocity to be consistent with the fixed position.

#### Step 1. Update the velocity

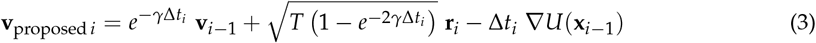

where the components of **r**_*i*_ follow a standard normal distribution.

In the limit for Δ*t*_*i*_ → 0, Equation (3) matches the rate of change of the mean and rate of added variance in equation (1). Equation (3) has also been carefully chosen to have stable behavior for large Δ*t* and/or *γ*. The first term decays the existing velocity by a factor of 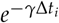. If Equation (1) only contained the first term, this would be the exact solution. This decay reduces the variance of the velocity by a factor of 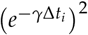. The second term re-injects variance sufficient to restore the variance of the velocity in every direction (orthogonal to constraints) as *T*.

A small refinement is made to avoid random rotation in the plane of projection, as this can be unsettling to view. Any part of the random noise **r**_*i*_ within the plane of the projection is subtracted out before the noise is added to the velocity. More precisely, considering the noise in matrix form **R**_*i*_ in the same way as the projection matrix **X**_*i* _−_ 1_, the projection of each row of **R**_*i*_ onto each row of **X**_*i* _−_ 1_ is subtracted from that row of **R**_*i*_. Previous tour software has also avoided this type of rotation, but in a different way, by using a “geodesic interpolation” method that operates between planes rather than frames (see Buja et al., 2005).

#### Step 2. Update the position based on the velocity

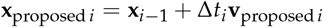

#### Step 3. Fix the updated position to be an orthonormal projection matrix

Now considering the proposed position vector as a projection matrix, we take the singular value decomposition and set the singular values to 1.

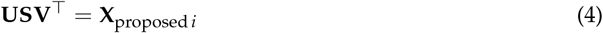

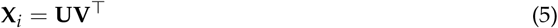

Here **U** is a 2 × 2 orthonormal matrix, **S** is a 2 × 2 diagonal matrix, and **V** is a *p* × 2 matrix with orthonormal columns. Let *s*_*j*_ be the values along the diagonal of **S**, the singular values, all of which are non-negative.

Equation (5) chooses the closest orthonormal projection matrix in terms of Euclidean distance to **x**_proposed *i*_. Stated another way, for proposed projection matrix **X**, this is the matrix **A** that minimizes the Frobenius norm ∥**X** − **A** ∥. To see this, consider first the problem of finding the nearest orthonormal projection matrix to **U**^⊤^**X**.

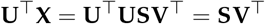

For each row in 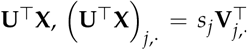, the nearest unit vector will be parallel to this vector, Namely 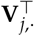,We know that the rows of **V**^⊤^ are orthogonal, so **V**^⊤^ is the nearest orthonormal projection matrix to **U**^⊤^**X**. Multiplying both of these matrices by an orthonormal matrix does not change the Frobenius norm of their difference, so the nearest orthonormal projection matrix to **UU**^⊤^**X** = **X** is **UV**^⊤^.

#### Step 4. Fix the velocity to be consistent with the fixed position

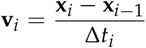

### Guiding projections using the potential energy function

We can use any function we like for the potential energy *U*(**x**), so long as we are able to calculate its gradient. This is used in **langevitour** both as a method of interaction and to provide a set of automatic guides.

When an energy function is being used, the temperature *T* plays a role analogous to variance in the normal distribution. When the temperature is very low, the system seeks the minimum of the energy function. As the temperature is raised, projections further and further from the minimum are produced.

### Linear energy function for interaction

For some choice of vector **a**, we can set the energy function to be the dot product

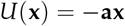

This encourages the projection to have a large component parallel to **a**. In **langevitour** this is used when labels are dragged onto the plot area to control the placement of particular axes of the high-dimensional space or to control the position of the mean of a group of points.

### Central force energy function

The Box-Cox power transformation (Box and Cox, 1964) provides a useful building block for energy functions.

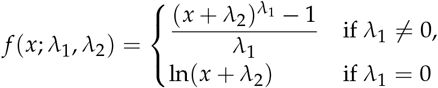

An energy function creating forces away from or toward the center can be defined using:

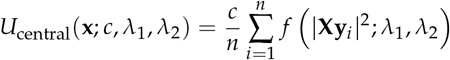

**langevitour** offers a central “push” guide (*c* < 0, *λ*_1_ = 0.5, *λ*_2_ = 0.0001) and a central “pull” guide (*c* > 0, *λ*_1_ = 0.5, *λ*_2_ = 0.0001). These cone-shaped energy functions result in a nearly constant magnitude outward or inward force on points, except for a small region close to the center.

### Layout by point-point repulsion

It was found that repulsion forces between pairs of points can be used to produce an informative layout. Let *S* be a set of pairs of points (*i, j*). Ideally we would make this the set of all possible pairs of points, but for computational efficiency **langevitour** uses a random mini-batch of 5,000 pairs of points per iteration, with a different mini-batch used each time. The use of random mini-batches to approximate the gradient injects extra noise into the system (see Mandt et al., 2017). The effect is similar to increasing the temperature slightly.

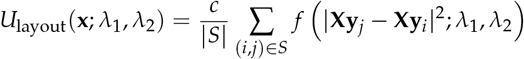

The power parameter *λ*_1_ determines whether the layout is governed by long range or short range forces. **langevitour** offers “ultra-local” (*c* < 0, *λ*_1_ = −1, *λ*_2_ = 0.0025), “local” (*c <* 0, *λ*_1_ = 0, *λ*_2_ = 0.0001), “PCA” (*c* < 0, *λ*_1_ = 1, *λ*_2_ = 0), and “outlier” (*c* < 0, *λ*_1_ = 2, *λ*_2_ = 0) guides. The “local” guide is the preferred default. With this guide, pairs of points that are near to each other exert more force than pairs of points that are far apart. The “ultra-local” guide potentially produces better layouts but is somewhat unstable. The “PCA” guide is equivalent to PCA. The “outlier” guide seeks projections where there are some points that are very far from other points.

### Blending energy functions

A sum of energy functions such as the above can be used to produce behavior that is a blend of the behaviors from the individual functions. For example, one or more labels could be dragged onto the plot, and also a guide activated.

### scRNA-Seq processing steps

Kang et al. (2018) processed sequencing reads using the 10x Genomics CellRanger software and provided the resulting count data in the Gene Expression Omnibus (GEO) database as accession number GSE96583. They also provided their annotation of the cells into different types, and their doublet detection based on genetic differences between individuals. There are 29,065 cells in total. 3,169 of these are identified as doublets.

Normalization by total count per cell, log transformation, and PCA were carried out using the **Seurat** package. As per Seurat defaults, only the top 2,000 highly variable genes are used.

Doublets containing cells from the same individual can not be identified by genetic differences, so Bioconductor package **scDblFinder** (Germain et al., 2022) was used to impute further doublets using the recoverDoublets function. This only works between cells of different types, but doublets containing cells of the same type are not really a problem. A further 595 doublets were identified this way.

The top PCs capture as much variation in the data as possible, but are not necessarily individually interpretable. To aid biological interpretation, it would be better if each component represented changes in the expression of a distinct set of genes. We would like each differentially expressed gene to have loadings that are mostly concentrated in a single component, and for convenience we would prefer for this loading to be positive. With these goals in mind, the varimax rotation of the gene loadings was found using the varimax() function in **stats**, with Kaiser normalization disabled. Both the gene loadings and the cell scores are rotated. Then, for each component, if the loadings have negative skew both the loadings and scores are negated.

To allow a smooth frame-rate in **langevitour**, the dataset was sub-sampled down to 10,000 cells.

### k-nearest neighbor denoising method

The denoising method implemented in the function knnDenoise() is as follows. We first find the k-nearest neighbors to each point (including the point itself). We then update each point to be the average of the set of points reachable within a certain number of steps along the directed k-nearest neighbors graph. Here *k* = 30 and two steps were used.

This method is loosely inspired by the k-nearest neighbors smoothing method of Wagner et al. (2018), as well as the use of the nearest neighbor graph in UMAP.

### Source code

The R code used to process the scRNA-Seq data is given in the supplemental file processing.R. Code for figure generation from the processed data is given in figures.R. Example code to control the widget using HTML buttons is given in the R Markdown document figures-page.Rmd.

## Discussion

Langevin Dynamics produces samples from a specified probability distribution, and produces a path suitable for animation. A possible future application would be to use this to visualize uncertainty in a Bayesian statistical model, while placing details of the model directly under interactive control. Langevin Dynamics is similar to the Hamiltonian Monte Carlo method (HMC, see Neal, 2011) used for example by Stan (Carpenter et al., 2017). Usually HMC alternates steps of completely randomizing the velocity and relatively long runs of Hamiltonian Dynamics simulation. This would produce a continuous path, but with occasional sharp turns. However Neal (2011) describes a version of HMC with frequent partial velocity refreshment that closely resembles what has been used in this article (apart from the use of constraints). An accept/reject step can make the sampling precisely accurate even with discrete time-steps. Another possibility, for large datasets, is to use mini-batch gradients for computational efficiency (Mandt et al., 2017). Mini-batch gradients provide a noisy estimate of the full gradient, and with careful tuning the level of noise required for Langevin Dynamics will be injected into the velocity.

Motion provides a channel of visual information not possible in static images. We are not accustomed to visualizing objects in more than three dimensions, but things that move together in the natural environment are usually physically connected, and this seems to be how our eyes interpret the small rotations in more than three dimensions displayed by **langevitour**.

A Javascript widget has been introduced for interactively exploring high-dimensional data. It is readily usable from the R environment, or in Shiny websites, or in HTML documents generated using R Markdown, including static HTML reports, slideshows, and journal articles.

## Supporting information

R code and interactive HTML figures

## Acknowledgements

The idea for **langevitour** arose during discussions with Prof. Di Cook and Dr. Stuart Lee at Monash University.

