## Supplementary material for "langevitour: smooth interactive touring of high dimensions, demonstrated with scRNA-Seq data": R code and interactive HTML figures: figures-page.html


### Figures

#### Figure 1

`palmerpenguins` data visualized using
`langevitour`.

The interface allows:

- Setting a “guide.” This causes to pursue projections in the
  vicinity of the minimum of an energy function. For example the PCA guide
  seeks projections with large variance in both the x and y directions.
  PCA
  None
- Hiding particular groups by unchecking their checkbox, in order
  to focus on other groups. For example by hiding Gentoo penguins we can
  focus on the difference between Adelie and Chinstrap penguins. The guide
  is also only applied to the visible groups.
  Hide
  Gentoo
  Show
  all
- Hiding particular axes by unchecking their checkbox. For example
  without bill length Adelie and Chinstrap penguins can no longer be
  distinguished.
  Hide
  bill length
  Show
  all
- Dragging labels on to the plot to concentrate on particular axes
  or try to separate out a particular group. The projection may not
  exactly match the label positions, since it must still be orthonormal.
  Bill
  length
  Bill
  length and depth
  None
- Adjusting damping. High damping produces jerky Brownian motion.
  Less damping produces smoother less random motion. The fastest way to
  thoroughly explore the space of projections is an intermediate damping
  level.
  Brownian
  Smooth
  Default
- Adjusting heat. More heat makes the projection move faster, and
  also strays further from the optimum projection defined by a guide or
  any labels that have been dragged on to the plot.
  Hot
  Cool
  Default
- Mousing over a group label. The group is highlighted.
- Mousing over an axis label. A scale and rug are displayed, and
  points are colored according to their position on that axis.

#### Figure 2

scRNA-Seq analysis using Seurat. (A) Scree plot. (B) UMAP layout
based on the cell scores of the first 10 PCs. U are unstimulated and S
are stimulated cells.

#### Figure 3

scRNA-Seq cell scores visualized using langevitour.

This is a moderately complex dataset. It may be helpful to initially
hide
the stimulated cells and
activate
the “local” guide so as to see the relationships between
unstimulated PBMC cells. The guided layout may not be able to perfectly
separate all the clusters as UMAP can, but small motions make clear
where clusters are overlapping by accident rather than real proximity.
For example B cells and T cells are distinct.
Hiding
the doublets makes the clusters more distinct.
Show
all

#### Figure 4

Noise reduction of scRNA-Seq cell scores.

#### Figure 5

Noise reduced scRNA-Seq cell scores.

Doublets still lie between clusters in this denoised version. Compare
this to UMAP layout, where the doublets tend to be attached to one or
other cluster.
Hiding
the doublets again makes clusters cleaner.
Show
all

Components associated with particular cell types can be found.
C1
for monocytes,
C5
for CD8 T cells and NK cells, and
C6
for B cells. The response to the cytokine is mostly contained in
C3
with a further monocyte specific response in
C2.
Reset

#### Figure 6

scRNA-Seq gene loadings visualized using langevitour.

It can be seen that biological phenomena such as different cell types
(C5,
C6)
or response to the cytokine
(C3)
often involve distinct sets of genes. Furthermore, cell types are
defined more by the activation of genes than the deactivation of genes.
Varimax rotation was able to align these distinct sets of genes into
specific components. An exception is genes used by monocytes and the
monocyte response to stimulation
(C1
and C2). These two components show a broad fan of genes, which
can be interpreted as the genes involved in being a monocyte also being
involved to varying degrees in the monocyte response to stimulation.
Reset
