## Supplementary material for "langevitour: smooth interactive touring of high dimensions, demonstrated with scRNA-Seq data": R code and interactive HTML figures: figures-page.Rmd

````
---
title: Figures
---

<style type="text/css">
.main-container {
  max-width: 100% !important;
  margin: auto;
}
</style>

```{r echo=FALSE}
knitr::opts_chunk$set(echo = FALSE, warning = FALSE, message = FALSE)
```

```{r}
source("figures.R")


# Function to display a button

go <- function(label, widget, state) {
    command <- paste0(
        "document.getElementById(\"",widget,"\")",
        ".langevitour.setState(",state,");",
        "document.getElementById(\"",widget,"\")",
        ".scrollIntoView({behavior:\"smooth\"})")
    command <- htmltools::htmlEscape(command, attribute=TRUE)

    knitr::asis_output(paste0(
        "<button onclick=\"",command,"\">",
        label,
        "</button>"))
}
```

# {.tabset}

## Figure 1

```{r}
figurePenguins$width <- "100%"
figurePenguins
```

`palmerpenguins` data visualized using `langevitour`.

The \CRANpkg{langevitour} interface allows: 

* Setting a "guide." This causes \CRANpkg{langevitour} to pursue projections in the vicinity of the minimum of an energy function. For example the PCA guide seeks projections with large variance in both the x and y directions.
`r go("PCA","penguins","{pointRepulsionType:'pca'}")`
`r go("None","penguins","{pointRepulsionType:'none'}")`

* Hiding particular groups by unchecking their checkbox, in order to focus on other groups. For example by hiding Gentoo penguins we can focus on the difference between Adelie and Chinstrap penguins. The guide is also only applied to the visible groups.
`r go("Hide Gentoo","penguins","{guideType:'pca',labelInactive:['Gentoo']}")`
`r go("Show all","penguins","{guideType:'pca',labelInactive:[]}")`

* Hiding particular axes by unchecking their checkbox. For example without bill length Adelie and Chinstrap penguins can no longer be distinguished.
`r go("Hide bill length","penguins","{labelInactive:['bill_length_mm']}")`
`r go("Show all","penguins","{labelInactive:[]}")`

* Dragging labels on to the plot to concentrate on particular axes or try to separate out a particular group. The projection may not exactly match the label positions, since it must still be orthonormal.
`r go("Bill length","penguins","{guideType:'none',labelPos:{bill_length_mm:[0.0,0.9]},labelInactive:[]}")`
`r go("Bill length and depth","penguins","{labelPos:{pointRepulsionType:'none',bill_length_mm:[0.0,0.5],bill_depth_mm:[0.5,0.0]},labelInactive:[]}")`
`r go("None","penguins","{labelPos:{}}")`

* Adjusting damping. High damping produces jerky Brownian motion. Less damping produces smoother less random motion. The fastest way to thoroughly explore the space of projections is an intermediate damping level.
`r go("Brownian","penguins","{damping:2}")`
`r go("Smooth","penguins","{damping:-2}")`
`r go("Default","penguins","{damping:0}")`

* Adjusting heat. More heat makes the projection move faster, and also strays further from the optimum projection defined by a guide or any labels that have been dragged on to the plot.
`r go("Hot","penguins","{heat:3}")`
`r go("Cool","penguins","{heat:-3}")`
`r go("Default","penguins","{heat:0}")`

* Mousing over a group label. The group is highlighted.

* Mousing over an axis label. A scale and rug are displayed, and points are colored according to their position on that axis.


## Figure 2

```{r}
figureStandard
```

scRNA-Seq analysis using Seurat. (A) Scree plot. (B) UMAP layout based on the cell scores of the first 10 PCs. U are unstimulated and S are stimulated cells.


## Figure 3

```{r}
figureRaw$width <- "100%"
figureRaw
```

scRNA-Seq cell scores visualized using langevitour.

This is a moderately complex dataset. It may be helpful to initially 
`r go("hide the stimulated cells", "raw", "{labelInactive:['S CD8 T cell','S Monocyte','S CD4 T cell','S Doublet',
'S B cell','S Other','S NK cell']}")` 
and 
`r go('activate the "local" guide', "raw", "{guideType:'local'}")` 
so as to see the relationships between unstimulated PBMC cells. The guided layout may not be able to perfectly separate all the clusters as UMAP can, but small motions make clear where clusters are overlapping by accident rather than real proximity. For example B cells and T cells are distinct. `r go("Hiding the doublets", "raw", "{labelInactive:['U Doublet','S Doublet']}")` makes the clusters more distinct. `r go("Show all","raw","{labelInactive:[]}")`


## Figure 4

```{r}
figureDenoising$width <- "100%"
figureDenoising$height <- NULL
figureDenoising
```

Noise reduction of scRNA-Seq cell scores.


## Figure 5

```{r}
figureDenoised$width <- "100%"
figureDenoised
```

Noise reduced scRNA-Seq cell scores.

Doublets still lie between clusters in this denoised version. Compare this to UMAP layout, where the doublets tend to be attached to one or other cluster.
`r go("Hiding the doublets", "denoised", "{labelInactive:['U Doublet','S Doublet']}")` again makes clusters cleaner.
`r go("Show all", "denoised", "{labelInactive:[]}")`

Components associated with particular cell types can be found.
`r go("C1","denoised","{labelPos:{C1:[0,0.9]}}")` for monocytes, 
`r go("C5","denoised","{labelPos:{C5:[0,0.9]}}")` 
for CD8 T cells and NK cells, and 
`r go("C6","denoised","{labelPos:{C6:[0,0.9]}}")`
for B cells.
The response to the cytokine is mostly contained in
`r go("C3","denoised","{labelPos:{C3:[0,0.9]}}")`
with a further monocyte specific response in
`r go("C2","denoised","{labelPos:{C3:[0,0.9],C2:[0.9,0]}}")`.
`r go("Reset","denoised","{labelPos:{}}")`


## Figure 6

```{r}
figureGene$width <- "100%"
figureGene
```

scRNA-Seq gene loadings visualized using langevitour.

It can be seen that biological phenomena such as different cell types 
(`r go("C5","gene","{labelPos:{C5:[0,0.9]}}")`, 
`r go("C6","gene","{labelPos:{C6:[0,0.9]}}")`) 
or response to the cytokine 
(`r go("C3","gene","{labelPos:{C3:[0,0.9]}}")`) 
often involve distinct sets of genes. Furthermore, cell types are defined more by the activation of genes than the deactivation of genes. Varimax rotation was able to align these distinct sets of genes into specific components. An exception is genes used by monocytes and the monocyte response to stimulation 
(`r go("C1 and C2","gene","{labelPos:{C1:[0.9,0],C2:[0,0.9]}}")`). 
These two components show a broad fan of genes, which can be interpreted as the genes involved in being a monocyte also being involved to varying degrees in the monocyte response to stimulation.
`r go("Reset","gene","{labelPos:{}}")`
````
