## Supplementary material for "langevitour: smooth interactive touring of high dimensions, demonstrated with scRNA-Seq data": R code and interactive HTML figures: figures.R

```
##
## Step through this script to generate the interactive figures in the article.
##   OR
## Look at figures-page.html (generated from figures-page.Rmd).
## 
## This script uses the file out.rds generated by processing.R.
##
## You will need to have installed some packages:
#
# install.packages(c("langevitour", "tidyverse", "cowplot", "palmerpenguins"))
#

library(langevitour)
library(tidyverse)
library(cowplot)
library(palmerpenguins)


## Palmer Penguins

completePenguins <- na.omit(penguins[,c(1,3,4,5,6)])
scale <- apply(completePenguins[,-1], 2, sd)*4

figurePenguins <- langevitour(
    completePenguins[,-1], 
    completePenguins$species, 
    scale=scale, pointSize=2, 
    elementId="penguins")

figurePenguins


## scRNA-Seq figures

out <- readRDS("out.rds")

stimLabel <- factor(ifelse(out$stim=="stim","S","U"), c("U","S")) 
crossLabel <- fct_cross(stimLabel, out$typeLabel, sep=" ")

h <- c(0,2,1,3,4,5,0)/6*360
c <- c(1,1,1,1,1,1,0)*100
colsCrossLabel <- hcl(h=rep(h,each=2), c=rep(c,each=2), l=rep(c(40,70),7), fixup=TRUE)
colsTypeLabel <- hcl(h=h, c=c, l=50, fixup=TRUE)


ggplot() + aes(x=out$umapX, y=out$umapY, color=crossLabel) + geom_point(stroke=0,size=1) + coord_fixed() + scale_color_discrete(type=colsCrossLabel) + theme_void()

subfigureScree <- ggplot() + 
    aes(x=1:30, y=out$stdev[1:30]**2) +
    geom_line() + 
    geom_point(size=1) + 
    labs(x="PCA component", y="Variance of component") + 
    scale_x_continuous(breaks=c(5,10,15,20,25)) +
    scale_y_continuous(breaks=c(1,40,80)) +
    coord_cartesian(xlim=c(0,31), ylim=c(0,80), expand=FALSE) +
    theme_cowplot(font_size=11) 

subfigureUmap <- ggplot() + 
    aes(x=out$umapX,y=out$umapY,color=crossLabel) + 
    geom_point(size=0.75, stroke=0) + 
    scale_color_discrete(type=colsCrossLabel) +
    scale_x_continuous(breaks=c()) +
    scale_y_continuous(breaks=c()) +
    guides(color = guide_legend(override.aes = list(size=4))) +
    coord_fixed() +
    labs(x="",y="",color="",title="") +
    theme_void() + 
    theme(legend.key.height = unit(1,"lines"))

figureStandard <- plot_grid(
    subfigureScree, subfigureUmap, 
    labels="AUTO", label_fontface="plain", label_size=10, 
    rel_widths=c(1,2), align="h")

figureStandard


## langevitour widget of the cell scores

figureRaw <- langevitour(
    out$vmScores, 
    crossLabel, 
    levelColors=colsCrossLabel, 
    elementId="raw")

figureRaw


## langevitour widget illustrating denoising

nTake <- 1000
denoisingLabel <- fct_cross(
    rep(stimLabel[1:nTake], 2), 
    rep(fct_inorder(c("Original","Denoised")), each=nTake), 
    sep=" ")
denoisingCol <- hcl(h=c(60,60,250,250), c=c(140,140,140,140), l=c(50,70,50,70), fixup=TRUE)

figureDenoising <- langevitour(
    rbind(out$vmScores[1:nTake,], out$denoisedScores[1:nTake,]),
    denoisingLabel,
    levelColors = denoisingCol,
    lineFrom = 1:nTake,
    lineTo = (1:nTake)+nTake,
    state = '{"axesOn":false,"guideType":"local"}',
    pointSize = 1.25,
    height = "500px",
    elementId="denoising")

figureDenoising


## langevitour widget of the denoised cell scores

figureDenoised <- langevitour(
    out$denoisedScores, 
    crossLabel, 
    levelColors=colsCrossLabel,
    state = '{"guideType":"local"}',
    elementId="denoised")

figureDenoised


## langevitour widget of the gene loadings

figureGene <- langevitour(
    out$vmLoadings, 
    name=rownames(out$vmLoadings), 
    scale=0.45, 
    elementId="gene")

figureGene
```
