## Supplementary material for "langevitour: smooth interactive touring of high dimensions, demonstrated with scRNA-Seq data": R code and interactive HTML figures: processing.R

```
##
## Run this script to download and process data.
##
## You will first need to install some packages:
#
# install.packages(c("langevitour", "tidyverse", "Seurat", "BiocManager"))
# BiocManager::install("scDblFinder")
#
## Then run the script:
#
# source("processing.R")
#

library(langevitour)
library(tidyverse)
library(Seurat)
library(scDblFinder)

set.seed(1)

# Download data from Gene Expression Omnibus (GEO) ====

dir.create("data", showWarnings=FALSE)

# Download from GEO can be slow.
options(timeout=3600)

if (!file.exists("data/GSE96583_RAW.tar") || 
    file.info("data/GSE96583_RAW.tar")$size != 76195840) {
    download.file(
        "https://www.ncbi.nlm.nih.gov/geo/download/?acc=GSE96583&format=file",
        "data/GSE96583_RAW.tar")
}

if (!file.exists("data/GSE96583_batch2.total.tsne.df.tsv.gz") ||
    file.info("data/GSE96583_batch2.total.tsne.df.tsv.gz")$size != 756342) {
    download.file(
        "https://ftp.ncbi.nlm.nih.gov/geo/series/GSE96nnn/GSE96583/suppl/GSE96583_batch2.total.tsne.df.tsv.gz",
        "data/GSE96583_batch2.total.tsne.df.tsv.gz")
}

if (!file.exists("data/GSE96583_batch2.genes.tsv.gz") ||
    file.info("data/GSE96583_batch2.genes.tsv.gz")$size != 277054) {
    download.file(
        "https://ftp.ncbi.nlm.nih.gov/geo/series/GSE96nnn/GSE96583/suppl/GSE96583_batch2.genes.tsv.gz",
        "data/GSE96583_batch2.genes.tsv.gz")
}


untar("data/GSE96583_RAW.tar", exdir="data")


# Load the data ====

cells <- read.table("data/GSE96583_batch2.total.tsne.df.tsv.gz", sep="\t")

counts <- cbind(
    ReadMtx("data/GSM2560248_2.1.mtx.gz", "data/GSM2560248_barcodes.tsv.gz", "data/GSE96583_batch2.genes.tsv.gz"),
    ReadMtx("data/GSM2560249_2.2.mtx.gz", "data/GSM2560249_barcodes.tsv.gz", "data/GSE96583_batch2.genes.tsv.gz"))

colnames(counts) <- rownames(cells)


# Seurat processing ====

obj <- CreateSeuratObject(counts, meta.data=cells)
obj <- NormalizeData(obj)
obj <- FindVariableFeatures(obj, selection.method = "vst", nfeatures = 2000)
obj <- ScaleData(obj)
obj <- RunPCA(obj, "RNA")
obj <- RunUMAP(obj, dims=1:10)


## Varimax rotation ====

nDim <- 10

loadings <- obj$pca@feature.loadings[,1:nDim]
scores <- obj$pca@cell.embeddings[,1:nDim]

# Find varimax rotation (without Kaiser normalization)
rotation <- varimax(loadings, normalize=FALSE)$rotmat

# Tidy the rotation: flip components to have positive skew
flips <- ifelse(colSums((loadings %*% rotation) ** 3) < 0, -1, 1)
rotation <- sweep(rotation, 2, flips, '*')

# Tidy the rotation: order by variance of scores
rotation <- rotation[, rev(order(colSums((scores %*% rotation)**2)))]

colnames(rotation) <- paste0("C",1:nDim)

vmLoadings <- loadings %*% rotation
vmScores <- scores %*% rotation


## Denoising ====

denoisedScores <- knnDenoise(vmScores, k=30, steps=2)


## Doublet detection ====
#
# Some doublets have already been identified based on genetic differences between individuals.
#
# Attempt to identify further doublets where the two cells came from the same indiviudal. 
#
# Detection is done on the unstimulated and stimulated cells independently since there will be no cells that are a mixture of unstimulated and stimulated.
#

labelDoublets <- function(obj) {
    reco <- recoverDoublets(
        obj$pca@cell.embeddings, 
        doublets=obj$multiplets == "doublet", 
        samples=table(obj$ind),    # Relative proportions of cells from each sample, here individual donors
        transposed=TRUE)

    case_when(
        reco$known ~ "known",
        reco$predicted ~ "predicted",
        TRUE ~ "singlet")
}

doubletLabel <-
    split(seq_len(ncol(obj)), obj$stim) |> 
    map(~labelDoublets(obj[,.])) |> 
    unsplit(obj$stim)


## Simplified labels ====

typeLabel <- case_when(
        doubletLabel != "singlet" ~ "Doublet",
        obj$cell == "B cells" ~ "B cell",
        obj$cell == "CD4 T cells" ~ "CD4 T cell",
        obj$cell == "CD8 T cells" ~ "CD8 T cell",
        obj$cell == "NK cells" ~ "NK cell",
        obj$cell == "CD14+ Monocytes" ~ "Monocyte",
        obj$cell == "FCGR3A+ Monocytes" ~ "Monocyte",
        TRUE ~ "Other"
    ) |> factor(c(
        "Doublet",
        "CD4 T cell",
        "CD8 T cell",
        "NK cell",
        "B cell",
        "Monocyte",
        "Other"
    ))


## Write out results ====
#
# Only 10,000 cells are kept out of 29,065, so langevitour can run at a smooth framerate.
#

keep <- sample.int(ncol(obj), 10000)

saveRDS(file="out.rds", object=list(
    stim = obj$stim[keep],
    vmLoadings = vmLoadings,
    vmScores = vmScores[keep,],
    denoisedScores = denoisedScores[keep,],
    doubletLabel = doubletLabel[keep],
    typeLabel = typeLabel[keep],
    umapX = as.vector(obj$umap@cell.embeddings[keep,1]),
    umapY = as.vector(obj$umap@cell.embeddings[keep,2]),
    stdev = Stdev(obj)))
```
